# HCCDB v2.0: Decompose the Expression Variations by Single-cell RNA-seq and Spatial Transcriptomics in HCC

**DOI:** 10.1101/2023.06.15.545045

**Authors:** Ziming Jiang, Yanhong Wu, Yuxin Miao, Kaige Deng, Fan Yang, Shuhuan Xu, Yupeng Wang, Renke You, Lei Zhang, Yuhan Fan, Wenbo Guo, Qiuyu Lian, Lei Chen, Xuegong Zhang, Yongchang Zheng, Jin Gu

## Abstract

Large-scale transcriptomic data are crucial for understanding the molecular features of hepatocellular carcinoma (HCC). By integrating 15 transcriptomic datasets of HCC clinical samples, the first version of HCCDB was released in 2018. The meta-analysis of differentially expressed genes and prognosis-related genes across multiple datasets provides a systematic view of the altered biological processes and the inter-patient heterogeneities of HCC with high reproducibility and robustness. After four years, the database needs to integrate recently published datasets. Furthermore, the latest single-cell and spatial transcriptomics provided a great opportunity to decipher the complex gene expression variations at the cellular level with spatial architecture. Here, we present HCCDB v2.0, an updated version that combines bulk, single-cell, and spatial transcriptomic data of HCC clinical samples. It dramatically expands the bulk sample size, adding 1656 new samples of 11 datasets to the existing 3917 samples, thereby enhancing the reliability of transcriptomic meta-analysis. A total of 182,832 cells and 69,352 spatial spots are added to the single-cell and spatial transcriptomics sections, respectively. A novel single-cell level and 2-dimension (sc-2D) metric was proposed to summarize the cell type-specific and dysregulated gene expression patterns. Results are all graphically visualized in our online portal, allowing users to easily retrieve data through a user-friendly interface and navigate between different views. With extensive clinical phenotypes and transcriptomic data in the database, we show two applications for identifying prognosis-associated cells and tumor microenvironment. HCCDB v2.0 is available at http://lifeome.net/database/hccdb2.

## Introduction

Hepatocellular carcinoma (HCC), which accounts for the vast majority (75%–85%) of primary liver cancer, is one of the leading digestive system malignancies [1]. The accumulation of transcriptomic data in HCC has facilitated the precise subtyping and biomarker identification [2–4]. However, due to the high heterogeneity of HCC, the transcriptomic data from a single cohort frequently generate inconsistent results due to limited sample size. The meta-analysis is an important approach to identify the stable patterns and the cohort-specific effects across different datasets [5]. To provide such a resource for studying the heterogeneities and the dysregulated biological processes in HCC, we have developed HCCDB (v1.0) which integrated transcriptomes of 3917 samples from 15 bulk datasets and emphasized the centrality of meta-analysis in transcriptomic analysis [6–8]. Due to the growth of published datasets of HCC clinical samples over the past four years, it’s imperative to increase the volume of the database contents. Moreover, it is interesting to assess the stability or reproducibility of the meta-analysis results after adding new datasets.

The bulk transcriptomic data provide important resources for analysing the gene expression variations of tumors in terms of malignancy, aggressiveness, and cell composition. However, bulk data only provide the average gene expressions of the sample which consists of multiple cell types and tumor sub-clones. The latest single-cell RNA sequencing (scRNA-seq) and spatial transcriptomics (ST) can decompose gene expression variations at the cellular level and obtain the spatial distribution of intact tissues. The scRNA-seq technique captures the cellular heterogeneity within the same tissue and reveals distinct cell subpopulations [9,10]. ST preserves the spatial location information of tumor tissues by in situ characterization of tissue spots, shedding light integrating the functional and structural aspects of transcriptomic analysis [11]. Emerging techniques have made it possible to identify the dominant cell populations and spatial patterns of gene expression variations. However, the high cost of these techniques limits their use to large clinical cohorts. The integration of the strengths of scRNA-seq and ST, along with the valuable clinical information from traditional bulk transcriptomics studies, holds great promise for providing a comprehensive transcriptional landscape of HCC.

To provide a unified portal for studying the gene expression variations in HCC with both large population and single-cell resolution, we released HCCDB v2.0, an updated version containing 5573 bulk transcriptomic samples, 182,832 cells, and 69,352 spatial spots. Extra to the previous 4D metric for bulk data, a single-cell level and 2-dimension (sc-2D) metric for single-cell data was also proposed for depicting the gene expression variations across different cell types and the differential expressions between tumor and normal hepatocytes. Based on this resource, we successfully identified cell subpopulations and tumor microenvironments associated with poor prognosis by integrating the three-way transcriptomic data. To facilitate the usage of this resource, a new searchable web portal was designed to visualize a set of pre-calculated results.

## Database content and computation methods

### The archived bulk expression datasets

For the release of HCCDB 2.0, we collected publicly accessible expression datasets of HCC up to March 2022. In total, 24 HCC datasets and 5572 samples were archived (Table S1). The preprocessing procedures were described in our previous article. Log2 transformation was applied to probe values from microarray and normalized read counts from RNA sequencing, respectively. Clinical information, such as tumor stages and survival time, was also collected if available. We standardized the terminology of clinical information so that it could be analyzed across datasets. For instance, the unit of all survival time information was converted to months.

### Search and processing of single-cell RNA sequencing datasets

We first retrieved scRNA-seq datasets of both human normal liver and HCC tissue in Gene Expression Omnibus (GEO) database. By only considering datasets with 3’-end sequencing from 10x Genomics platform, two scRNA-seq datasets were archived, including a normal liver dataset (HCCDB-SC1) [12] and a HCC dataset (HCCDB-SC2) [13]. In order to enrich the datasets in the database, we enrolled 7 HCC patients, and obtained 7 tumor samples and 3 tumor-adjacent liver tissue (HCCDB-SC3, in-house data). The information of single-cell transcriptomic datasets was shown in Table S2. We carried out cell-level quality control by filtering mitochondrial mRNA and unique feature counts [14]. Cells with elevated mitochondrial mRNA levels (mt > 20) were excluded, as well as those with excessively high or low unique feature counts (fc < 200 or fc > 10,000). Principal Components Analysis (PCA) was carried out to project the spots onto a low-dimensional space defined by the first 30 principal components (PCs). We eliminated batch effects between datasets through the CCA algorithm. The clustering was performed using Seurat (v4.3.0) [15]. Two dimensionality reduction methods, UMAP and TSNE, were used to visualize the clustering. We manually annotated the cell clusters and identified 7 major lineages and 18 minor lineages in total.

### Processing of spatial transcriptomics

We collected the ST data from our prior article [16]. For ST, we released 17 tissue sections with a total of 69,352 tissue spots from 5 patients (HCC-1 to HCC-5). The detailed processing method was described as before. In total, four nontumor sections, four leading-edge sections, four tumor sections, one portal vein tumor thrombus section, and four sections from an intact tumor nodule were obtained (Table S3).

### Identification of driver factors regulating gene expression

DNA methylation and somatic copy number variations (CNV) were both considered as major factors affecting gene expression. To determine genetic factors that contribute to the regulation of transcript expression, we applied a multivariate linear regression model to DNA methylation and CNV data from The Cancer Genome Atlas (TCGA) datasets. Considering Gene i, CpG methylation levels in the promoter region (M), the somatic copy number variations (C), and the random error (ω), the equation was characterized as follows:

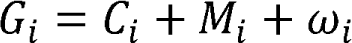

Individual genetic factors with adjustment *P* value < 0.05 (Benjamini-Hochberg correction) were considered as driver regulation factors.

### Identification of consistently differentially expressed genes

In HCCDB v2.0, the number of datasets with both tumor and adjacent samples was increased to 19. We used the t-test function in R on each dataset to determine if there was a significant difference in gene expression between tumor and adjacent samples, followed by Benjamini-Hochberg correction. Consistently differentially expressed genes (cDEGs) were defined as those with expression measurements in 8 datasets and significant differential expression (adjusted *P* < 0.001 and |log2 fold change| > 0.6) in at least half of the datasets.

### Prognostic analysis

Six datasets (HCCDB6, HCCDB15, HCCDB18, HCCDB19, HCCDB24, and HCCDB25) with overall survival time information were used to evaluate the prognostic performance of each gene. The median value of the genes in each dataset was employed to classify the high and low expression groups. The significance was determined by the log-rank test. Prognostic genes were defined as genes with adjusted *P*L<L0.001 (Benjamini-Hochberg correction) in one dataset or adjusted *P*L<L0.01 in more than two datasets. Genes with negative Cox coefficients were labelled as “favorable genes”, indicating a higher expression value correlated with a lower risk. Genes with positive Cox coefficients were labelled as “unfavorable genes”.

### Definition of a 4D metric, sc-2D metric, and spatial transcriptomics deregulation metric

In the last release, we introduced a 4D metric for each gene, a potent tool to evaluate gene variation and summarize the expression pattern in bulk level, including liver-specific metric, deregulation metric, tumor-specific metric, and HCC-specific metric. Because of the increasing number of HCCDB datasets, the deregulation metric was revised. The definition of 4D metric was characterized as previously [6]. In addition, we proposed a sc-2D metric in single-cell scales, which conducted the two-dimensional metric for each gene in each cell type to decipher gene expression variance. Because of limited adjacent samples, we considered cells from both adjacent and normal tissues as the control group of HCC samples. Here are the definitions for the two metrics:

1. Cell-specific metric quantifies the specificity of the gene i in each cell type j in comparison with other cell types:

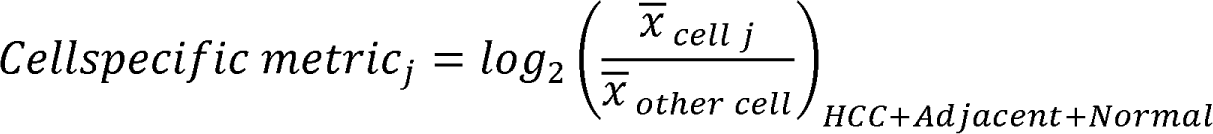
2. HCC deregulation metric quantifies the log_2_ fold change of gene i in tumor tissues compare with normal and adjacent tissues in each cell type j:

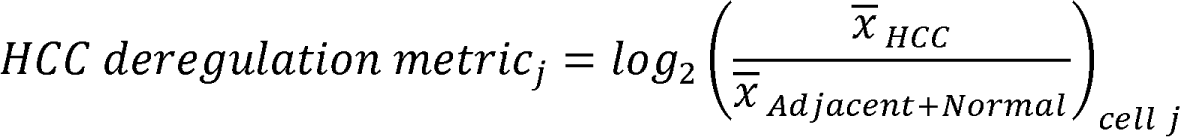

For ST sub-atlas, the ST deregulation metric was calculated as below:

1. ST deregulation metric quantifies the log2 fold change of gene i in tumor spots compare with adjacent spots in ST samples (HCC 1–5):

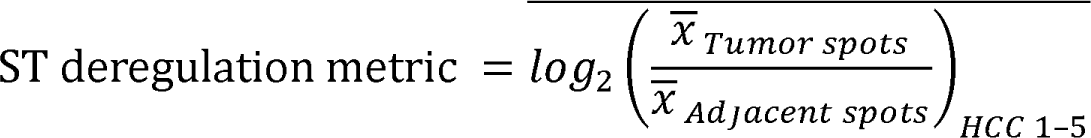

### Calculation of the highly regional genes score

We used the highly regional genes (HRG) algorithm to evaluate the regional distribution extent of individual genes [17]. The scoring function for gene g is defined as:

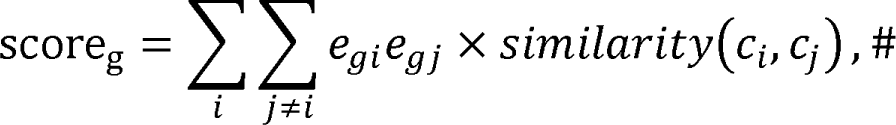

where *e_gi_* is scaled gene expression of gene g in cell c_i_ and *similarity*(*c_i_, c_j_*) is the spatial distance between cell c_i_ and *c_j_*. *similarity*(*c_i_, c_j_*) equals 1 If cell *c_i_* and *c_j_* are connected in space and similarity (*c_i_, c_j_*), otherwise. *e_gi_* is positive if the expression is higher than the average gene expression, and negative otherwise. If the expressions of gene g in cell *c_i_* and *c_j_* both higher or lower than the average expression level, *e_gi_e_gj_* × *similarity*(*c_i_, c_j_*) will positively contribute to its score. If *c_i_* and *c_j_* to be similar, then the contribution will be greater. On the contrary, if the expressions of gene g in *c_i_* and *c_j_* are either higher or lower than the average expression level, *e_gi_e_gj_* × *similarity*(*c_i_*, *c_j_*) will exert negative contribution to its score. Intuitively, high score indicates that expression levels in two similar cells are just similar so genes have regional distribution patterns.

The open-source HighlyRegionalGenes R package can be accessed on GitHub (https://github.com/JulieBaker1/HighlyRegionalGenes). We conducted 10 rounds of iteration and selected the top 2000 genes per round. The HRG score was log2 transformed and scaled accordingly.

### Identification of phenotype-related cells by Scissor

We applied Scissor to identify cell populations that are related to overall survival. Scissor is formulated as the following network regularized sparse regression model:

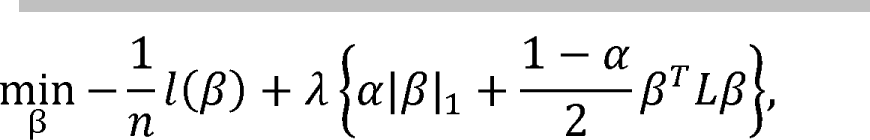

where L is a symmetric normalized Laplacian matrix, which is defined as

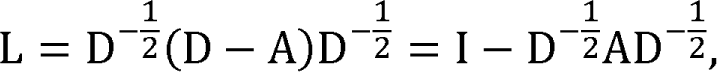

where *A* = (*a_ij_*)_m×m_ is a binary or weighted adjacency matrix of G. *a_ij_* equals one or a value ranging from 0 to 1 if cell *i*, and *j* are connected in G and *a_ij_* = 0, otherwise. *D* = (d_*ij*_)_m×m_ is the degree matrix of *G*, where *d_ii_* ∑_*m j*=1_, *a_ij_*, and d_*ij*_ = 0 for *i* ≠ *j*. The tuning parameter λ controls the overall strength of the penalty, and α balances the amount of regularization for smoothness and sparsity. n is the number of bulk sample number. β denote a vector of coefficients on cells and *l*(β) denote an appropriately chosen log-likelihood function.

For clinical survival data, Cox regression was employed to determine the most phenotype-associated cell subpopulations from the single-cell data.

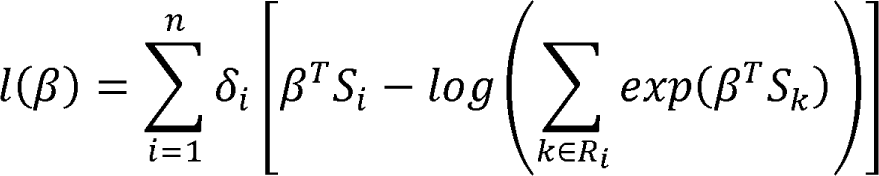

Where δ_*i*_ is the event indicator, *S_i_* = (*s*_*i*1_, *s*_*i*2_, …, *s*_*i*m_)^*T*^ as the correlation coefficients for sample, across all m cells.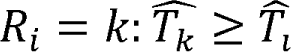 denotes the risk set at time *T̂*_*i*_.

The non-zero coefficients of f solved by the above optimization model are used (positive sign of f) cells are associated with worse survival, and Scissor− (negative to select the cell subpopulations associated with the overall survival. Scissor+ sign of f) cells are associated with good survival. A reliability significance test is further designed to control false associations. The final single cell overall survival status is merged by the survival status calculated from all the seven sets of bulk datasets.

The parameter alpha, which balances the impact of the l1 norm and network-based penalties, was set to 0.05. The cutoff value for the percentage of scissor-selected cells among all cells was set to 0.2.

### Estimate cell abundance of tumor microenvironment

To quantify the abundance of stromal and immune cells of data in HCCDB, we applied Xcell (R package, version 1.1.0) on the normalized expression data of each dataset to estimate scores for 38 infiltrating cell subtypes [18]. Kaplan-Meier analysis and log-rank test (R survival package, version 3.4-0) were applied to assess the clinical relevance of CD8^+^ T cells.

### Implementation and results

#### Overview of updated HCCDB database

The database has been expanded both horizontally and longitudinally. Horizontally, new bulk transcriptomic, scRNA-seq, and ST datasets have been added. Longitudinally, a new analysis pipeline has been created to decipher bulk gene expression variations using scRNA-seq and ST, providing a cellular level resolution of gene expression patterns (**Figure 1**).

**Figure 1.**
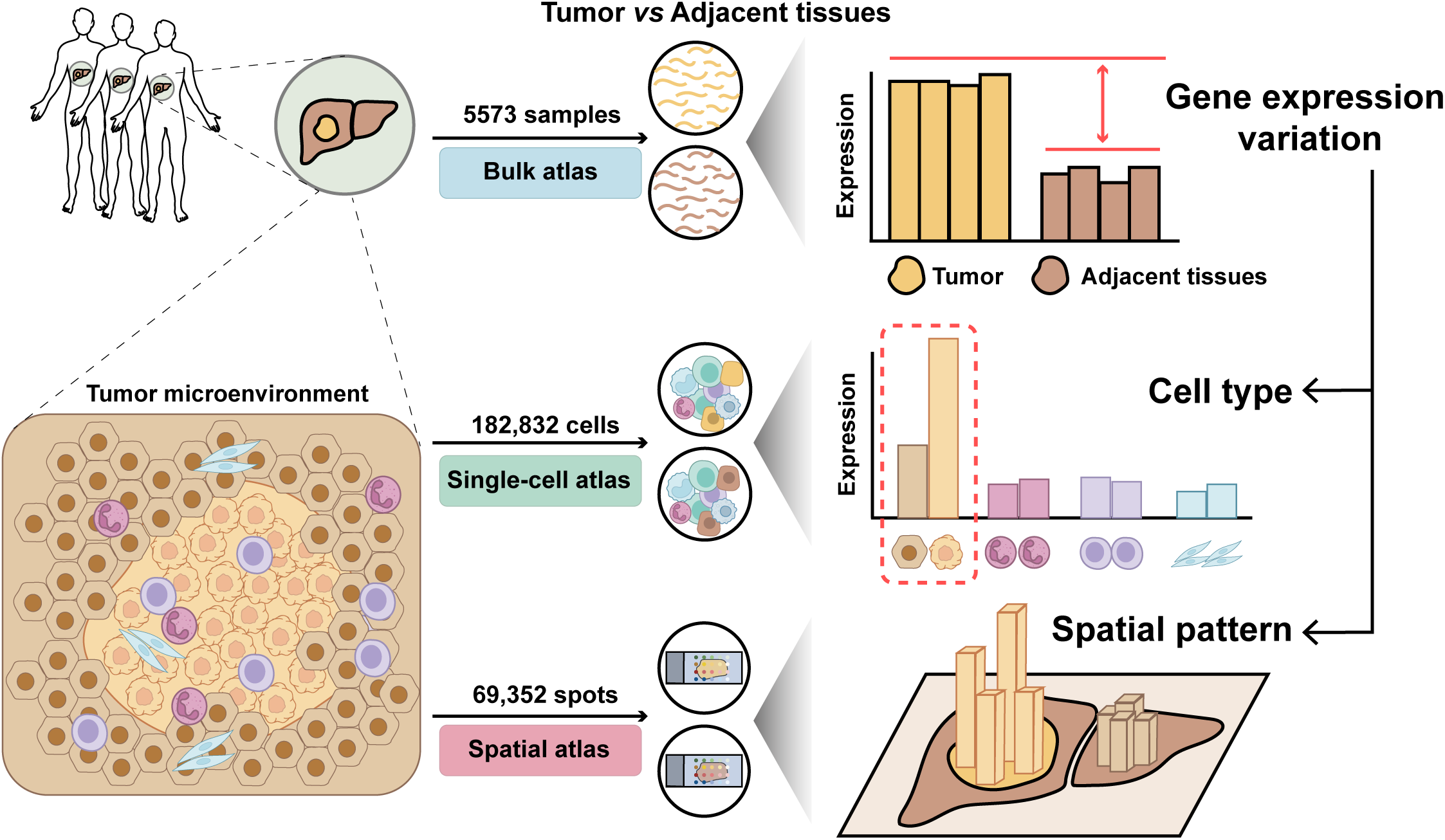
Overall design of HCCDB v2.0 HCCDB v2.0 offers transcriptomic data for both primary tumor tissues and adjacent liver tissues. It features three sub-atlases to give a comprehensive view of the transcriptomic landscape. The gene expression variations obtained through bulk transcriptomics can be further analyzed at the single-cell level through single-cell transcriptomics, providing a deeper understanding of the underlying mechanisms. Spatial transcriptomics, on the other hand, sheds light on the regional expression pattern and spatial architecture.

HCCDB v2.0 includes three sub-atlases: bulk transcriptomics, single-cell transcriptomics, and spatial transcriptomics, all of which are accessible from the HOME page. For queried genes, two search modes are available: single and multi-gene search. The result page of original single-gene search has been relocated to the bulk transcriptomics sub-atlas. The bulk atlas includes summary information, expression patterns, and survival analysis, while the co-expression panel has been removed to streamline content. Moreover, third-party links and PubMed database links have been reviewed and updated. The single-cell sub-atlas presents summary expression patterns using UMAP, violin plots, and dot plots. In ST sub-atlas, emphasis is placed on highlighting the advantages of ST in exploring tumor spatial heterogeneity. Hematoxylin and eosin (H&E) stain sections and point-to-point spatial gene expression distributions are provided. A parallel framework for the three sub-atlases allows users to switch between sub-atlases at every step of browsing, searching, and downloading.

#### Assessment of the meta-analysis stability for bulk transcriptomics after update

After finishing cleaning procedures, 1656 samples from 11 datasets were involved, which significantly increased the database content compared to the previous version, which had 3917 samples from 15 datasets. The new datasets include HCCDB19, HCCDB25, and HCCDB30 derived from RNA-seq platforms and the remaining datasets using microarrays. Seven of the eleven datasets contain both adjacent and HCC samples with standardized clinical information. The expanded datasets brought novel clinical phenotypes, including disease-free survival, tumor purity, and sorafenib response. In total, HCCDB v2.0 released 26 datasets with 5573 samples and 16 clinical phenotypes, 19 datasets of which have both adjacent and HCC samples (Table S1–S3). In addition, we attached a tag about genomic factors controlling gene expression to the result page of bulk transcriptomics. For example, the upregulation of *GPC3* in HCC was putatively driven by a methylation event of the promoter region.

We proposed metrics to classify the gene expression pattern of HCC, including the 4D metric, consistently differentially expressed genes, and prognostic genes, by integrating multiple datasets. We evaluated the impact of expanding the dataset size by 42% (1656/3917) on these three metrics. The revised deregulation metric showed a strong correlation with the original one (Table S4, Pearson correlation test, R^2^ = 0.90, *P* < 2.2×10^−16^). The top three upregulated genes, *GPC3*, *SPINK1*, and *AKR1B10*, remained the most significantly upregulated after the revision (**Figure 2**A). The number of cDEGs decreased from 1259 to 1065, with some changes in gene identity (Figure 2B), but no upregulated genes were converted to downregulated, and *vice versa*, showing the relative stability of cDEG identity. The number of prognostic genes increased from 1346 to 1893, indicating the discovery of more prognostic genes with the expansion of datasets (Figure 2C). In summary, our results demonstrate that gene expression pattern is essentially stable after expanding the datasets, but with added information and robustness.

**Figure 2.**
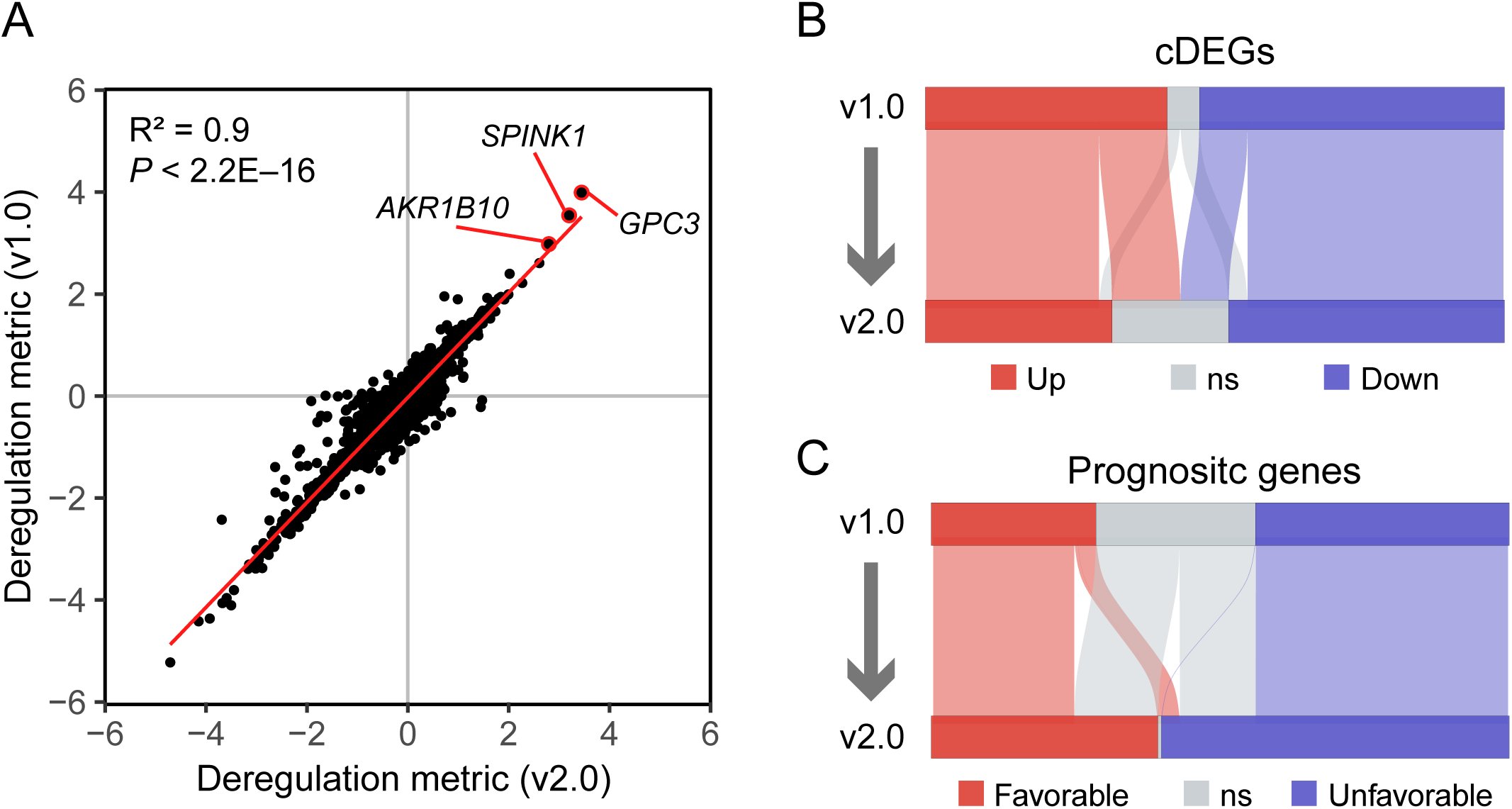
Analysis of bulk transcriptomic results after the expansion of datasets **A.** Scatter plot depicted the deregulation metric between the two versions (v1.0 for y axis and v2.0 for x axis). The red line represented the linear regression line. Top three genes with the highest deregulation metric in two versions were pinpointed. Statistical significance was assessed using Pearson correlation (*P* < 2.2×10^−16^). The alluvial plot showed changes in cDEGs (**B**) and prognostic genes (**C**) after the revision. cDEG, consistently differentially expressed gene.

#### Decipher the gene expression variations at single-cell level

To deepen the understanding of HCC’s transcriptional landscape, we added a scRNA-seq atlas featuring 182,832 cells from three datasets, HCCDB-SC1 (normal liver data) [12], HCCDB-SC2 (HCC data) [13], and HCCDB-SC3 (HCC and adjacent tissues, in-house data). The analysis identified 9 major and 19 minor cell lineages. We added a graphical interface to the scRNA-seq results page, providing users with selective information such as dataset, tissue, patient ID, Seurat clusters, and cell types. For each queried gene, four graphs displaying gene expression abundance are provided: a UMAP plot, a t-SNE plot, a violin plot, and a dot plot. These plots can be easily downloaded as high-quality graphics.

To measure the extent of tumor microenvironment deregulation in HCC, we developed the sc-2D metric, including the cell-specific metric and HCC deregulation metric, quantifying the cell specificity and cellular deregulation degree for individual genes (Table S5 and S6). As an example, the gene *PLVAP*, which is expressed by tumor endothelium [19,20], exhibited a notably high cell-specific metric and HCC deregulation metric in both the major and minor cell lineages (**Figure 3**A). Each gene was assigned a unique identity tag, with the cell type with the highest cell-specific metric referred to as the “master cell type”. Most of the genes (82.28%, 12,761/15,509) were primarily expressed in hepatocytes, malignant cells, endothelial cells, fibroblast cells, and NK/T cells (Figure 3B), demonstrating that these cell types mainly contributed to the bulk variations of HCC. Additionally, when focusing on prognostic genes and cDEGs from bulk transcriptomics, we found that 58.5% (556/950) of cDEGs and 44.4% (720/1620) of prognostic genes were primarily expressed in hepatocytes or malignant cells and exhibited significant differences in expression between normal and tumor cells, highlighting the crucial role of malignant cells in tumor dysregulation and prognosis (Figure 3C and D).

**Figure 3.**
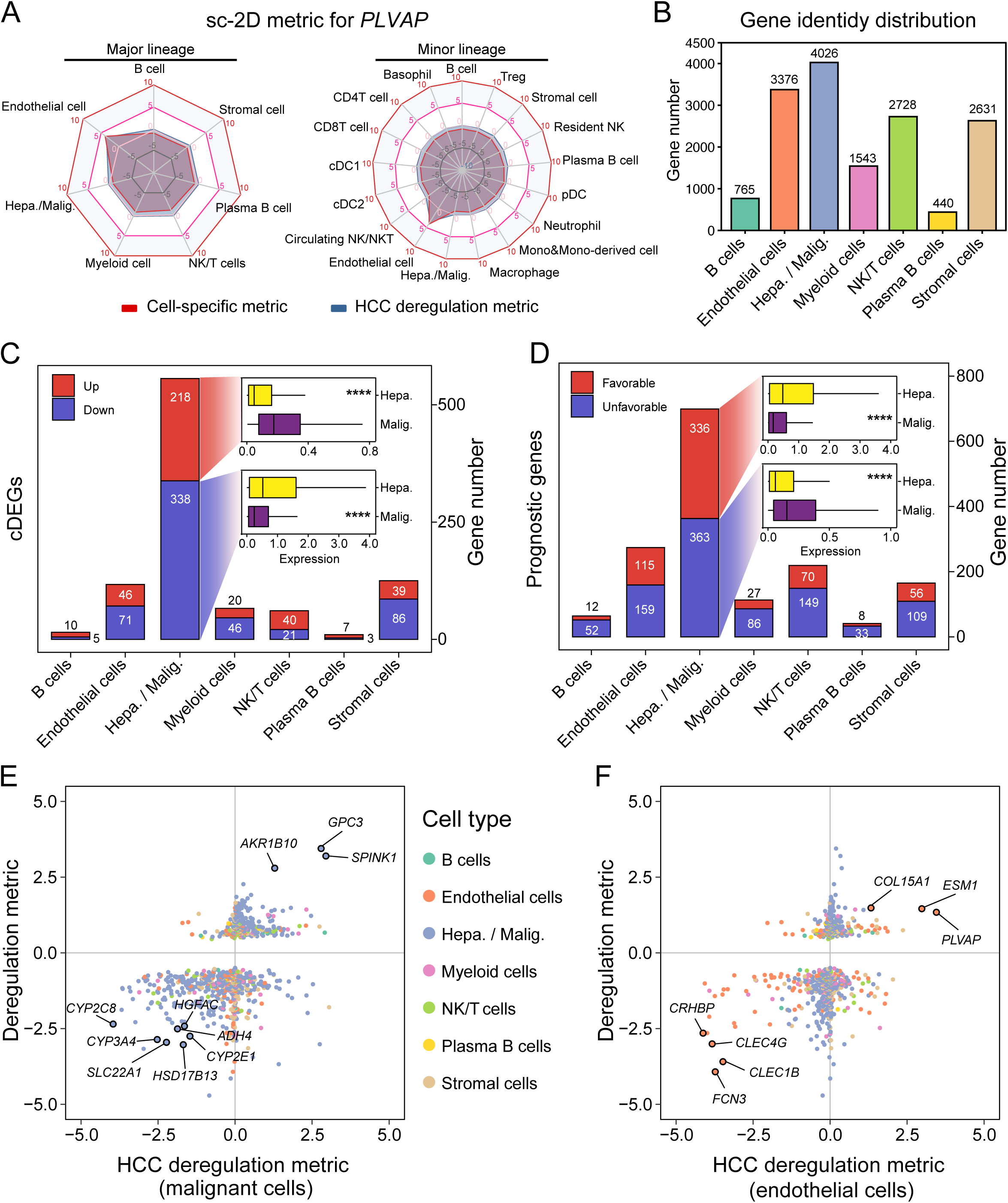
Analysis of sc-2D metric to decompose bulk gene expression patterns **A.** The radar plot illustrates the sc-2D pattern of an endothelial-specific gene, *PLVAP*, for the major lineage (left) and the minor panel (right). The cell-specific metric is depicted in red, while the HCC deregulation metric is depicted in blue. **B.** Distribution of gene identity. The cell type with the highest cell-specific metric was determined as cell-specific for individual genes. Distribution of gene identity for cDEGs (**C**) and prognostic genes (**D**), upregulated genes and favorable genes were colored by red; downregulated genes and unfavorable genes were colored by blue. The box chart showed the comparison of average expression of hepatocytes in normal liver tissues and malignant cells in tumor tissues. *P* value was calculated by the two-tailed Wilcoxon rank sum test. *, *P* < 0.05; **, *P* < 0.01; ***, *P* < 0.001, and ****, *P* < 0.0001. Scatter plots depicted the relationship between deregulation metric derived from bulk transcriptomics and HCC deregulation metric of malignant cells (**E**) and endothelial cells (**F**). Hepa., hepatocytes; Malig., malignant cells.

The deregulation degree of HCC was quantified by the deregulation metric derived from bulk transcriptomics. However, the contribution of each cell type to the deregulation of HCC is still unclear. Here, we deciphered the bulk deregulation metric by HCC deregulation metric of each cell type. *GPC3* and *SPINK1* were mainly expressed in malignant cells, with both high bulk deregulation metric and HCC deregulation metric of malignant cells (Figure 3E). Distinctive patterns of genes were shown in other cell types, such as up-regulated *THY1* in stromal cells and *SPP1* in myeloid cells (Figure S1A–E). Intriguingly, several genes, containing *CLEG4G* and *FCN3*, were expressed in normal endothelial cells but downregulated in tumor tissues (Figure 3F). *FCN3* has been identified as a tumor-suppressive gene in lung adenocarcinoma and is associated with good prognosis in HCCDB [21]. Our results suggested that endothelial-specific genes, such as *FCN3*, may play important roles in preventing tumor progression. However, the roles of endothelial cells and these endothelial-specific genes in HCC remain largely unexplored. Together, the expansion of single-cell atlas decomposed the bulk variations to the cellular level and identified potential therapeutic targets.

#### Spatial transcriptomics benefits the investigation of tumor heterogeneity

For ST atlas, we collected 69,352 tissue spots and 17 tissue histological sections from five patients (HCC-1 to HCC-5). The tissue regions were annotated and included immune, tumor, adjacent, and stromal. The H&E stain sections and spatial cluster distribution plots for each patient are presented in the ST atlas, along with interactive UMAP and t-SNE plots. To visualize gene expression in different patients and tissue regions, we provided spatial feature plots, dot plots, and violin plots. The comparison of ST deregulation metric with the bulk deregulation metric revealed a high concordance between ST and bulk transcriptomics (**Figure 4**A).

**Figure 4.**
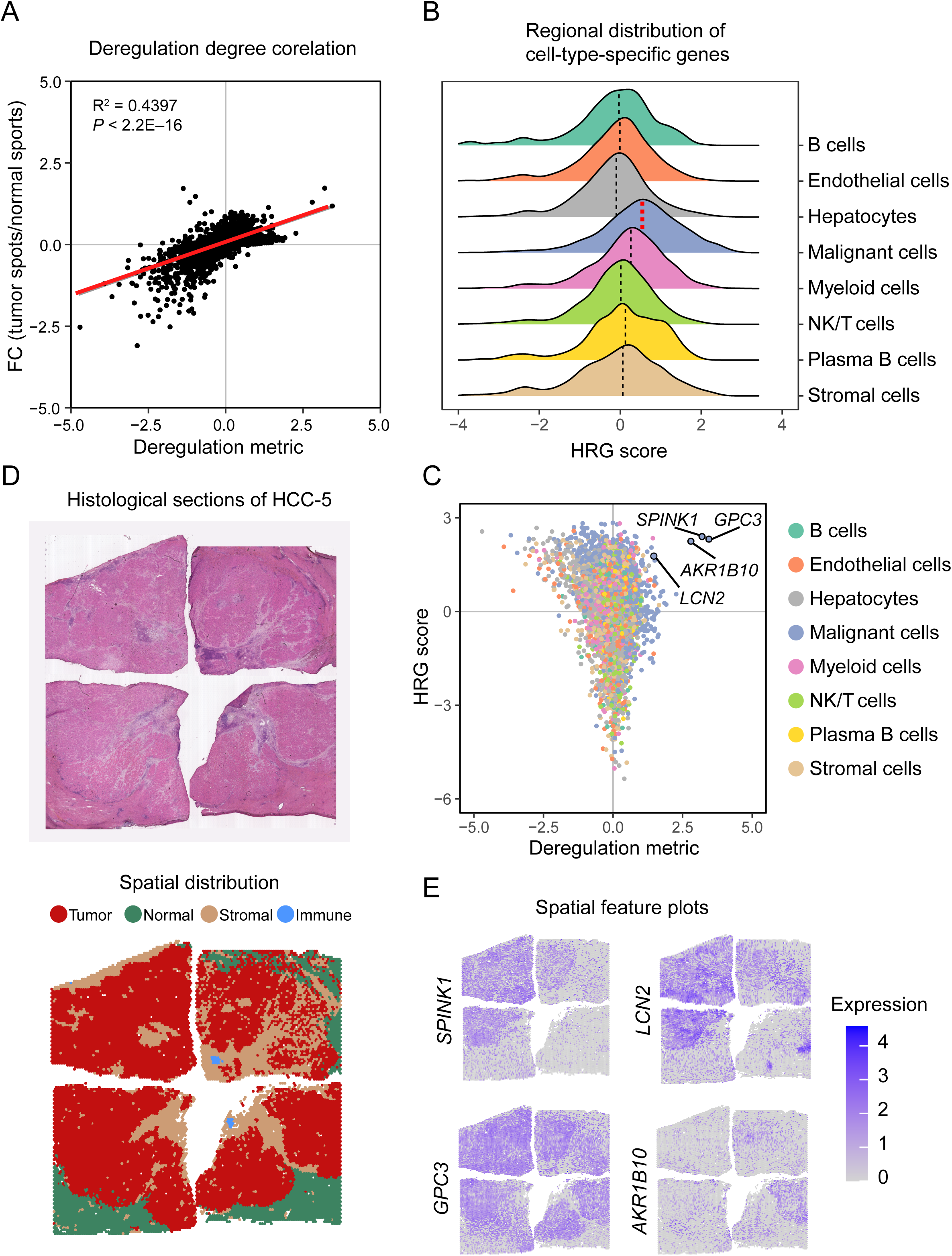
Spatial transcriptomics exhibits the intratumor heterogeneity **A.** Scatter plot depicted the correlation between deregulation metric derived from bulk transcriptomics (x axis) and ST deregulation metric derived from spatial transcriptomics (y axis), with the linear regression line represented in red. Pearson correlation was used to assess statistical significance, with a *P* value of less than 2.2×10^−16^. **B.** Regional distribution of cell-type-specific genes quantified by HRG score (see Methods). The median value for each group was indicated by dashed line. Malignant cell-specific genes showed the highest median HRG scores (colored by red). **C.** Scatter plot depicted the relationship between deregulation metric (x axis) derived from bulk transcriptomics and HRG score (y axis). Four genes with both high deregulation value and HRG score were pinpointed. **D.** H&E staining (top) of HCC-5 and the spatial cluster distribution (bottom). Four tissue spot types were identified including tumor (red), normal (green), stromal (yellow), and immune spots (blue). **E.** Spatial gene expression distribution of four genes (*GPC3*, *SPINK1*, *AKR1B10*, and *LCN2*) on HCC-5 (related to D). FC, fold change; ST, spatial transcriptomics; HRG, highly regional genes; H&E, hematoxylin and eosin.

To assess the ability of ST to detect intratumor heterogeneity, we utilized the HRG algorithm to analyze a 1-cm-diameter tumor nodule from patient HCC-5 [17]. The HRG algorithm, which identifies regionally distributed genes by constructing a cell-cell similarity graph, revealed that malignant-cell-specific genes determined by scRNA-seq had the highest HRG score among all cell types (Figure 4B). Additionally, several dysregulated genes, such as *GPC3*, *AKR1B10*, *SPINK1*, and *LCN2*, showed both high HRG scores and deregulation degree (Figure 4C, Table S7). Figure 4D and 4E shows different spatial expression distributions of these genes in this tumor nodule (Figure 4D and E). Even though these genes were extensively upregulated at the bulk level in different clinical stages, the spatial distribution patterns at the early developmental stages can be diverse. In conclusion, the combination of ST and bulk transcriptomic data improved our understanding of tumor heterogeneity and further interpretation of gene expression variations.

#### A population of prognosis-related cells identified by combining bulk transcriptomic and scRNA-seq dataset

HCCs are heterogeneous and include subpopulations such as cancer stem cells, which are known to drive tumor progression and poor prognosis [22]. To identify cells related to poor survival, we applied the Scissor method [23]. There are eight bulk datasets with survival information. We left out HCCDB15 which was derived from TCGA as the validation dataset. To avoid batch effects among bulk datasets, we transferred the survival information of the other bulk datasets to HCCDB-SC2 one by one. For those cells with contradicted transferred phenotype in different bulk datasets, we named them “uncertain”. Those cells related with good survival or poor survival in more than one bulk dataset were designated as good survival or poor survival cells respectively. Of the scissor-selected cells, most of them were malignant epithelial cells (**Figure 5**A and B, Figure S2A), indicating that epithelial cells are the decisive factors in patients’ survival. Besides, the phenotype of scissor-selected cells within a patient was consistent, either related to good survival or poor survival (Figure S2B). To identify key factors associated with worse survival, we compared poor survival cells to good survival cells (Figure 5C, Table S8).

**Figure 5.**
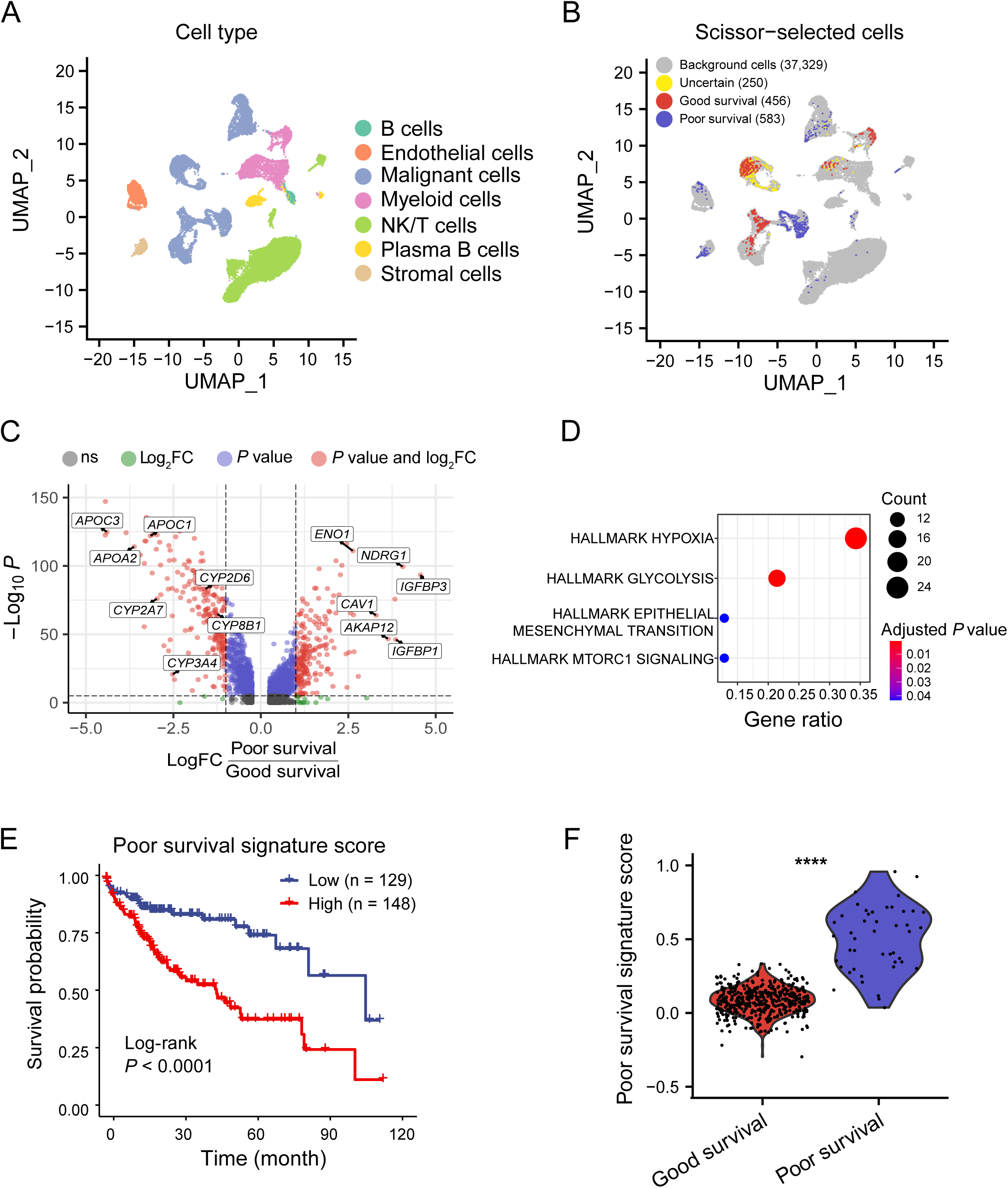
Prognosis-related-cell identification on malignant cells guided by HCCDB survival outcomes **A.** The UMAP visualization of major cell types from HCCDB-SC2. **B.** UMAP visualization of the scissor-selected cells. The red and blue dots are poor survival and good survival cells, respectively. The green dots are uncertain cells with contradicted transferred phenotype in different bulk datasets. **C.** Volcano plot of differential gene expressions in good survival cells versus poor survival cells. **D.** Enrichment bar plot of poor survival cells upregulated genes in Hallmark pathways. **E.** Kaplan-Meier survival curves show the clinical relevance of the poor survival signature on HCCDB15 dataset. **F.** The poor survival signature scores in HCCDB-SC3 scissor-selected cells. The FDR was the adjusted *P* value calculated by the two-tailed Wilcoxon rank sum test. *, *P* < 0.05; **, *P* < 0.01; ***, *P* < 0.001, and ****, *P* < 0.0001. FDR, false discovery rate.

Notably, we found hepatocyte function-related genes involved in glycogen/ lipid/ alcohol metabolism (APO/ALDH/ADH family genes), detoxification (CYP family genes), were relatively higher in good survival cells and multiple important hypoxia-related genes were highly expressed among poor survival cells (Figure S2C). Functional enrichment analysis also indicated that the poor-survival-related genes were enriched in the hypoxia pathway (Figure 5D). To validate the poor survival gene signature, we scored patients in validation datasets HCCDB15 and found that patients with higher scores had significantly poor survival (Figure 5E). At the same time, we applied the same procedure of choosing good survival and poor survival cells in HCCDB-SC3 dataset. We found that poor survival cells had significantly higher score of the poor survival gene signature derived from the HCCDB-SC2 dataset (Figure 5F), indicating the robustness of the poor survival signature. Taken together, the integration of scRNA-seq datasets and bulk datasets identified a specific population of prognosis-related cells and provided a poor survival signature for prognostic prediction.

#### Prognosis-related tumor microenvironment programs

Solid tumors are commonly infiltrated by immune cells which include T and B lymphocytes, natural killer (NK) cells, dendritic cells (DCs), macrophages, neutrophils, eosinophils, and mast cells [24]. To assess the relationship between tumor-infiltrating immune cells and tumor progression, we then applied Xcell on eight HCCDB datasets with phenotypic information to calculate the enrichment score of 38 types of immune cells and stromal cells (Table S9, see Methods). The Barcelona Clinic Liver Cancer (BCLC) system offers a prognostic stratification of HCC patients [25]. We observed that the enrichment scores of most immune cells decreased significantly with the deterioration of the disease in the four datasets with BCLC stage information (**Figure 6**A), which means that the abundance of immune infiltration also decreased with the deterioration of HCC. In the HCCDB28 dataset, we observed that the abundance of CD8^+^ T cells, DCs, eosinophils and NK cells in the samples that responded to sorafenib was significantly higher than that in the non-responsive samples. On the other hand, the abundance of other immune cells, such as CD4+ T cells, macrophages, monocytes, and neutrophils, was lower in the responder group (Figure 6B). Tumor mutational burden (TMB) is an important biomarker for response to PD-1/PD-L1 inhibitors [26]. We also explore the correlation between TMB and tumor-infiltrating immune cells using HCCDB19 dataset. The enrichment score of CD8^+^ T cells was positively associated with TMB significantly, whereas that of mast cells was negatively correlated (Figure 6C). Meanwhile, we found that the enrichment of CD8^+^ T cells was significantly associated with good survival in survival analyses (Figure 6D). Taken together, the integration of multiple datasets revealed that the tumor-infiltrating immune cells were closely related to clinical phenotype and tumor progression in HCC.

**Figure 6.**
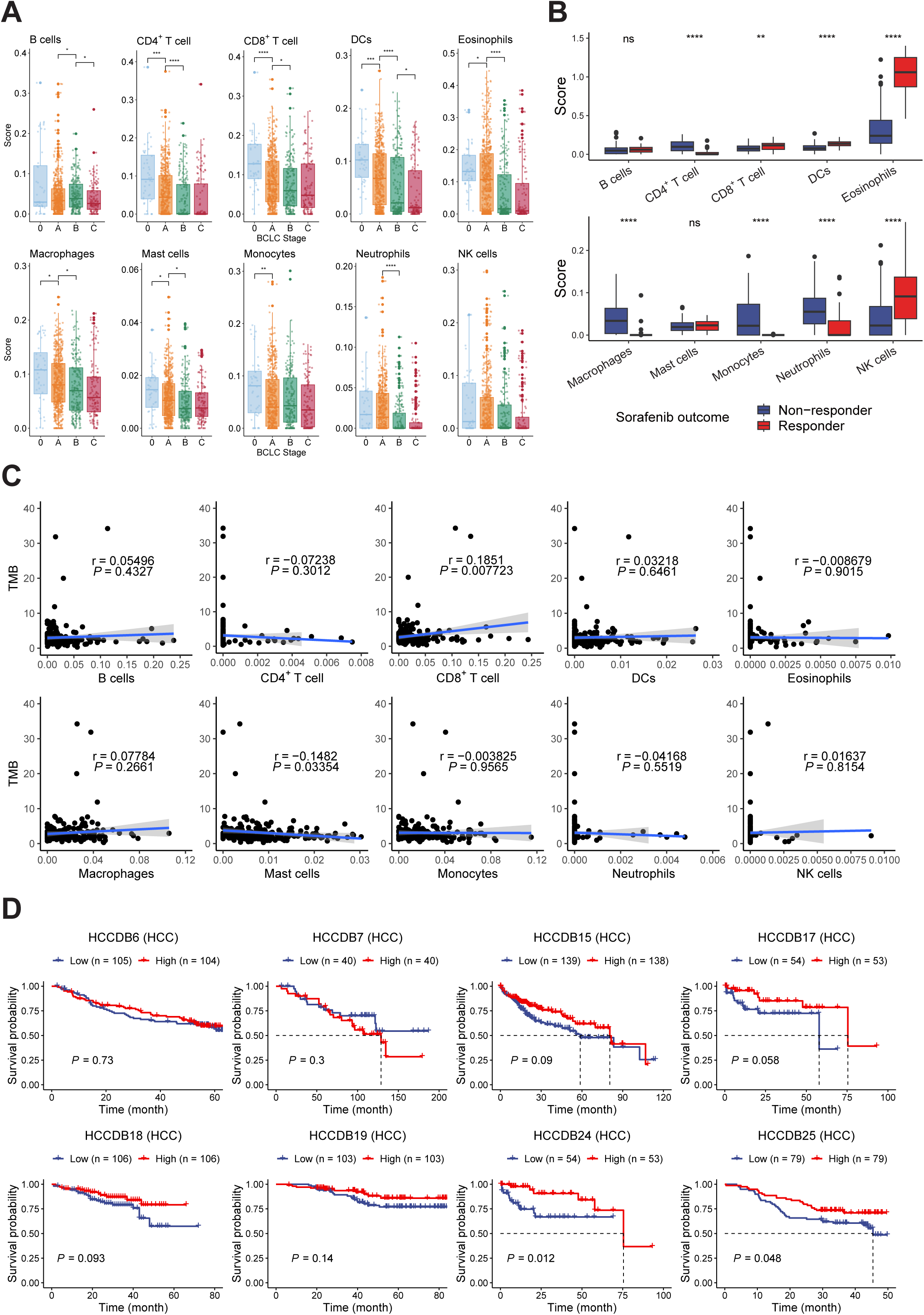
Deconvolution of HCC immune microenvironment **A.** Boxplot showing the connectivity between 10 tumor-infiltrating immune cells and the BCLC stage. The *P* value between the two groups is calculated with Wilcoxon tests. *, *P* < 0.05; **, *P* < 0.01; ***, *P* < 0.001, and ****, *P* < 0.0001. **B.** Boxplot showing the differences in the abundance of 10 immune cells between the sorafenib response group and the non-response group. The *P* value between the two groups is calculated with Wilcoxon tests. *, *P* < 0.05; **, *P* < 0.01; ***, *P* < 0.001, and ****, *P* < 0.0001. **C.** Scatter plots showing the relationship between the abundance of 10 tumor-infiltrating immune cells and TMB. Shaded bands represent 95% confidence intervals of linear regression slopes. *P* values are from t-tests. **D.** Kaplan-Meier plots showing the relationship between CD8^+^ T cells’ abundance and the clinical outcomes in HCC patients. The two groups of data were divided by the median of CD8^+^ T cell abundance. The statistical significance was determined by log-rank tests. BCLC, Barcelona Clinic Liver Cancer; TMB, tumor mutational burden; DCs, dendritic cells.

### Perspectives and concluding remarks

Large-scale transcriptomic data has facilitated the analysis and discovery of new therapeutic targets and biomarkers. Particularly, the rapid growth of bulk transcriptomic data provides a wealth of clinical information, which is advantageous for the integrative analysis of transcriptome and clinical information. However, the limited resolution of bulk transcriptomics hindered further analysis considering the growing need for the understanding of tumor heterogeneity and immune microenvironment. Therefore, we have released HCCDB v2.0, a comprehensive transcriptomic database with 5573 bulk transcriptomic samples, 182,832 cells, and 69,352 spatial spots. As far as we know, this is the first oncology database combining scRNA-seq and ST with bulk transcriptomics to describe transcriptional landscapes. To fully grasp the gene expression pattern, our parallel framework allows users to easily switch between different omics. Users can browse the comprehensive expression profiles of individual genes, including gene information and clinical information from the bulk transcriptomics atlas, cell annotation information from the scRNA-seq atlas, and spatial distribution from the spatial transcriptomics atlas.

Combined with genomic information and third-party links, users can easily achieve a one-stop searching workflow in HCCDB v2.0. 4D metric and sc-2D metric provide a new perspective for the integration analysis of different transcriptomic data, which is a bold and innovative attempt.

Here, we provide two cases for potential data exploration using the HCCDB v2.0 database. Scissor provides a good analytical model, which enables us to make full use of a large amount of clinical information of HCCDB v2.0 and transcriptomic data of scRNA-seq. By leveraging the abundant phenotype of the bulk datasets and the precision of single-cell datasets, we successfully identified phenotype-related cells. Scissor can transfer the phenotype to single cells. When transferring survival information in bulk datasets to HCCDB-SC2 scRNA-seq datasets, we identified epithelial cells related with poor survival whose upregulated genes are enriched in hypoxia hallmarks. These cells could provide potential drug targets for further investigation. The poor-survival signature can be successfully transferred to independent bulk datasets and scRNA-seq datasets, indicating the robustness of the signature. Xcell can identify the enrichment of multiple cell types in bulk data so that we can associate the clinical information in HCCDB v2.0 with one or several cell types. We observed lower immune infiltration correlated with elevated BCLC stage. In the HCCDB28 dataset, the abundance of different immune cells showed different distributions in sorafenib and control groups. Our data also indicated that CD8^+^ T cell enrichment significantly predicts high TMB and a good prognosis in HCC patients. When more phenotypes such as response to anti-PD1 therapy are collected in the future, we can identify cells related with the outcome of anti-PD1 therapy.

HCCDB v2.0 demonstrated the viability of data exploration to offer resources and convenience for resolving hot scientific issues. In the future, our group will keep the content updated, add new transcriptomic data, and integrate other omics data to create the new integrated analysis methodology. Thus, future iterations of HCCDB database may in fact demonstrate even greater potential.

## Data availability

HCCDB v2.0 is available at http://lifeome.net/database/hccdb2/.

## CRediT author statement

**Ziming Jiang**: Conceptualization, Methodology, Software, Formal analysis, Investigation, Data Curation, Writing - Original Draft, Visualization. **Yanhong Wu**: Formal analysis, Investigation, Data Curation, Writing - Original Draft, Visualization. **Yuxin Miao**: Formal analysis, Writing - Original Draft, Visualization. **Kaige Deng**, Investigation. **Fan Yang**, Investigation. **Shuhuan Xu**: Software. **Yupeng Wang**: Software. **Renke You**: Software. **Lei Zhang**: Software. **Yuhan Fan**, Investigation, Data Curation. **Wenbo Guo**, Methodology, Data Curation. **Qiuyu Lian**: Conceptualization, Methodology, Data Curation. **Lei Chen**, Conceptualization, Writing - Review & Editing. **Xuegong Zhang**, Conceptualization, Writing - Review & Editing. **Yongchang Zheng**, Conceptualization, Resources, Writing - Review & Editing, Funding acquisition. **Jin Gu**: Conceptualization, Methodology, Writing - Review & Editing, Supervision, Project administration, Funding acquisition. All authors read and approved the final manuscript.

## Acknowledgements

This work is funded by the National Key R&D Program of China (Grant No. 2021YFF1200901 awarded to JG), the National Natural Science Foundation of China (Grant Nos. 62133006, 61721003, and 62103273 awarded to JG), the Tsinghua University Initiative Scientific Research Program (Grant No. 20221080076 awarded to JG). Beijing Municipal Natural Science Foundation Project (Grant No. 7222130 awarded to YZ), Special Clinical Research Project of Peking Union Medical College Hospital (Grant No. 2022-PUMCH-A-236 awarded to YZ), CHEN XIAO-PING Foundation for the development of science and technology of HUBEI province (Grant No. CXPJJH1200008-10 awarded to YZ).

## Supplementary material

**Figure S1 Analysis of sc-2D metric in different cell types**

The scatter plot depicted the relationship between the deregulation metric derived from bulk transcriptomics and the HCC deregulation metric of stromal cells (**A**), myeloid cells (**B**), NK/T cells (**C**), B cells (**D**), and plasma B cells (**E**).

**Figure S2 Distribution of scissor-selected cells**

**A.** The constitution of scissor-selected cells in different cell types. **B.** The constitution of scissor-selected epithelial cells in different patients. **C.** Violin plots of expression levels of upregulated genes in good survival cells and poor survival cells. OS, overall survival.

**Table S1 Summary of bulk transcriptomics datasets**

**Table S2 Summary of single-cell transcriptomics datasets**

**Table S3 Summary of spatial transcriptomics datasets**

**Table S4 The revised 4D metrics**

**Table S5 Cell-specific metric of major cell lineages**

**Table S6 HCC deregulation metric of major cell lineages**

**Table S7 Scaled HRG score**

**Table S8 Poor survival markers**

**Table S9 Summary of tumor microenvironment information**

