## Supplementary figures and images for "HCCDB v2.0: Decompose the Expression Variations by Single-cell RNA-seq and Spatial Transcriptomics in HCC"

### Supplemental Figure 1

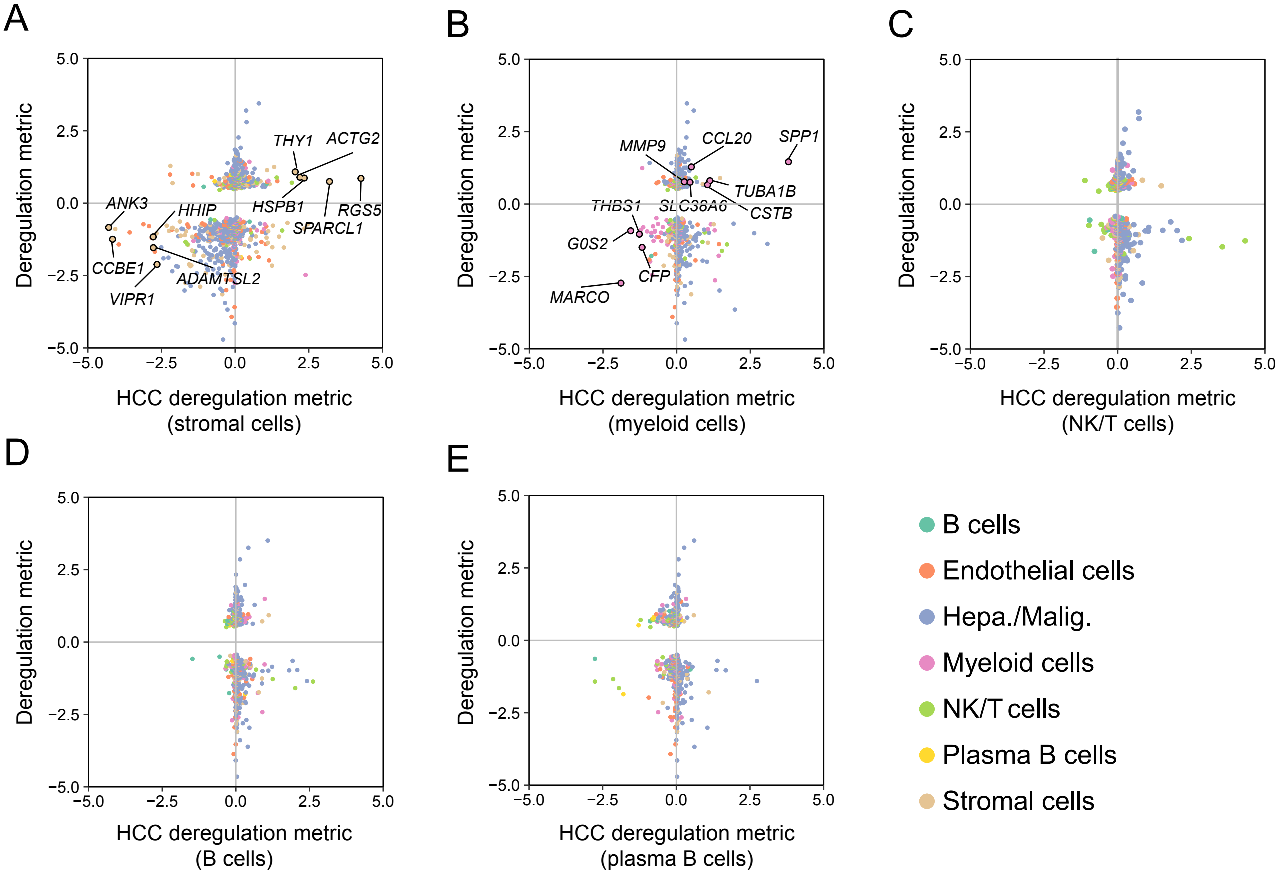

### Supplemental Figure 2

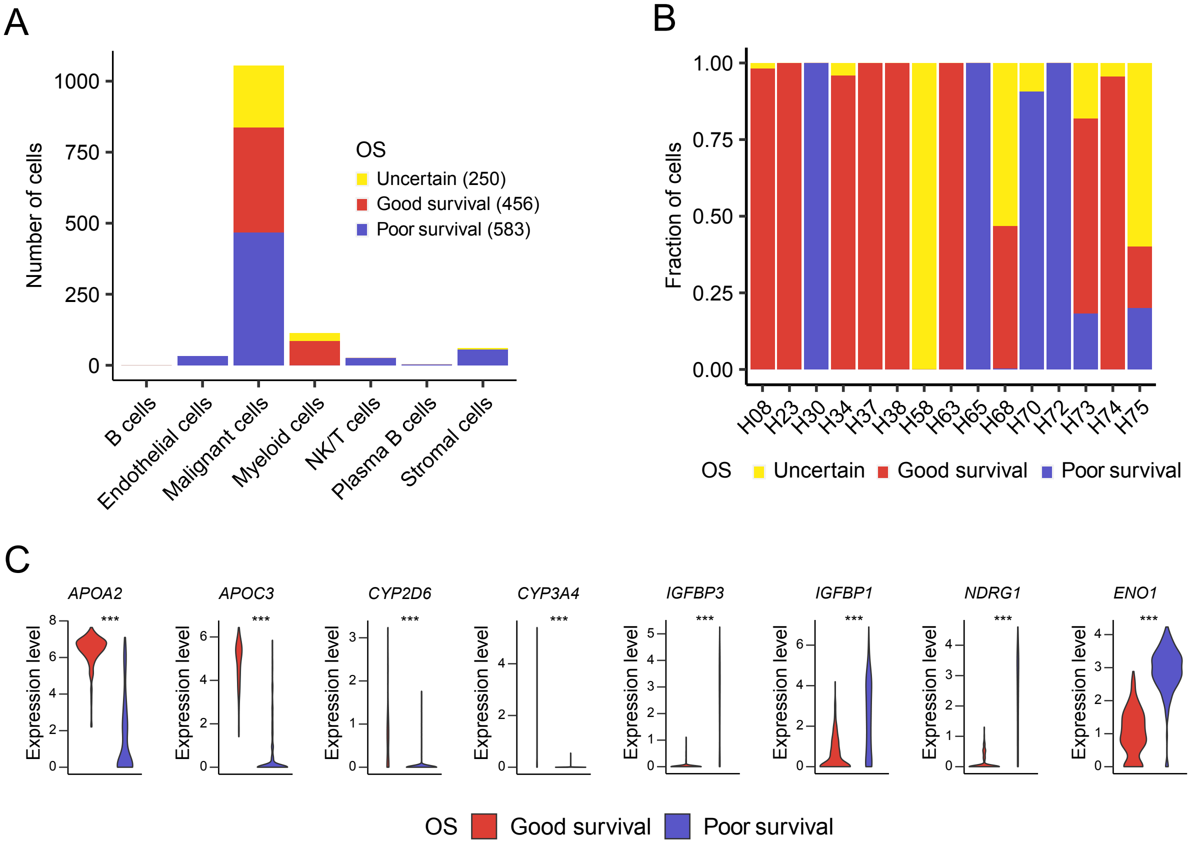
